# Frequency-dependent modulation of foveal contrast sensitivity by fine-scale exogenously triggered attention

**DOI:** 10.1101/2025.08.27.672541

**Authors:** Yue Guzhang, T. Florian Jaeger, Martina Poletti

**Affiliations:** Department of Brain and Cognitive Sciences, University of Rochester, Rochester, NY, United States; Center for Visual Science, University of Rochester, Rochester, NY, United States; Goergen Institute for Data Science and Artificial Intelligence, University of Rochester, Rochester, NY, United States; Department of Neurosciences, University of Rochester, Rochester, NY, United States

## Abstract

Exogenous attention is a rapid, involuntary mechanism that automatically reallocates processing resources toward salient stimuli. It enhances visual sensitivity in the vicinity of the salient stimulus, both in extrafoveal regions and within the high-acuity foveola. While the spatial frequencies modulated by exogenous attention in extrafoveal vision are well characterized, it remains unknown how this mechanism operates within the foveola, which can resolve spatial frequencies up to 30 cycles per degree (CPD). Here, we examined which spatial frequencies were enhanced by fine-grained deployments of exogenous attention within this highest-acuity region of the visual field. Using high-precision eye-tracking to precisely localize gaze during attentional allocation, we found that exogenous attention at the foveal scale selectively enhances contrast sensitivity for low- to mid-range spatial frequencies (4–8 CPD), with no significant benefits for higher spatial frequencies (12–20 CPD). In contrast, attention-related benefits on asymptotic performance at the highest contrast were observed across a wide range of spatial frequencies. These results indicate that, despite the high-resolution capacity of the foveola, exogenous attention remains an inflexible mechanism that, even at this scale, selectively enhances contrast gain for lower spatial frequencies—mirroring its behavior in extrafoveal vision.

## Introduction

Visual spatial attention is a fundamental mechanism that enables both humans (***Carrasco, 2011***) and animals (***Saban et al., 2017***; ***Hahner and Nieder, 2025***) to selectively process information from their environment. Often, shifts in spatial attention are accompanied by eye movements to focus on a specific location, a process known as overt spatial attention (***Kaspar, 2013***). However, covert spatial attention, the ability to shift attention independently of eye movements, is equally crucial in daily life. This ability enables us to monitor locations beyond our line of sight, such as when driving and keeping track of peripheral surroundings.

Covert spatial attention is typically categorized into two types: endogenous and exogenous attention. Endogenous attention refers to the voluntary allocation of processing resources to a specific location. While this shift occurs relatively slowly, taking approximately 200–300 ms to reach the target region, it can be sustained for an extended duration (***Theeuwes, 1994***; ***Findlay, 2003***; ***Chica and Lupiáñez, 2009***; ***Carrasco, 2011***; ***Chica et al., 2013***; ***Dugué et al., 2020***). In contrast, exogenous attention is driven by salient stimuli that automatically capture attention (***Theeuwes, 1994***; ***Findlay, 2003***; ***Chica and Lupiáñez, 2009***; ***Carrasco, 2011***; ***Chica et al., 2013***; ***Dugué et al., 2020***). This shift is rapid but transient, often followed by a phenomenon known as inhibition of return, moving attention away from the initially attended location (***Klein, 2000***; ***Chica and Lupiáñez, 2009***). Compared to endogenous attention, exogenous attention is more automatic and less flexible ***Corbetta and Shulman (2002***); ***Knudsen (2007***); ***Carrasco (2011***).

Until recently, research on the effects of covert attention on visual perception has focused primarily on extrafoveal vision. A vast body of literature has demonstrated that covert attention enhances visual contrast sensitivity (***Carrasco et al., 2000***; ***Martínez-Trujillo and Treue, 2002***; ***Reynolds and Chelazzi, 2004***; ***Pestilli and Carrasco, 2005***; ***Li et al., 2008***; ***Foster et al., 2021***) and increases spatial resolution (***Yeshurun and Carrasco, 1998***; ***Carrasco et al., 2002, 2006***; ***Jigo and Carrasco, 2018***) at selectively cued locations in the extrafovea. In contrast, attention within the high-acuity 1-deg foveola has often been considered uniform and distributed evenly throughout this small region. Therefore, the effects of attention in the fovea are traditionally studied using large stimuli encompassing one or more degrees of visual angle (***Miniussi et al., 2002***; ***Jigo and Carrasco, 2020***; ***Papaioannou and Luck, 2020***). However, recent findings showed that even within the 1-degree foveola, both endogenous (***Poletti et al., 2017***) and exogenous (***Guzhang et al., 2021***) attention can be covertly allocated in a highly spatially selective manner. For both types of covert attention, observers were better able to discriminate the orientation of fine details at an attended location—cued endogenously or exogenously—compared to nearby uncued locations just 0.26° away. Although these results highlighted the strikingly fine grain of attentional control, they also raised new questions. In particular, it remains unknown which spatial frequencies benefit from fine-grained attentional shifts within the foveola. While ***Guzhang et al. (2021***) demonstrated visual enhancement from exogenous attention at the foveal scale, the orientation discrimination task used in the study was relatively coarse, requiring observers to determine whether a stimulus was tilted ±45°. Despite the small size of the stimulus, such a task does not require high spatial frequencies (*e.g*., > 10 cycles per degree); in fact, frequencies around 4-8 cycles per degree (CPD) should be sufficient to perform it effectively. Therefore, the perceptual enhancement observed in ***Guzhang et al. (2021***) could be due to an enhancement of only lower or only higher spatial frequencies, or perhaps a broad range of spatial frequencies. The overall improvement in orientation discrimination of fine spatial stimuli is compatible with any of these scenarios.

The effects of *extra*foveal attention have been found to differ across spatial frequencies: while extrafoveal endogenous attention enhances a broad range of spatial frequencies (***Lu and Dosher, 2004***; ***Jigo and Carrasco, 2020***), extrafoveal exogenous attention selectively enhances high spatial frequencies (***Carrasco et al., 2006***; ***Barbot et al., 2011, 2012***; ***Carretié et al., 2012***; ***Jigo and Carrasco, 2020***; ***Fernández et al., 2022***), peaking just above the spatial frequency characterized by the highest sensitivity at a given eccentricity (***Jigo and Carrasco, 2020***). Whether fine spatial exogenous attention at the foveal scale modulates visual discrimination similarly is an open question. Generally, fine control of spatial attention at the foveal scale is required when examining fine spatial details, such as reading small text in a book or noticing subtle changes, like a traffic light switching or unexpected pedestrians from afar while driving (***Figure 1****A*). In these tasks, precise allocation of attention likely helps distinguishing and recognizing individual letters and details. It is possible that in the foveola, exogenous attention modulates a narrow range of spatial frequencies, similar to how it operates extrafoveally. However, while humans can resolve spatial frequencies up to 30 cycles per degree (CPD) in the foveola (***Intoy and Rucci, 2020***; ***Clark et al., 2022***), extrafoveally, spatial frequencies above 10 CPD cannot be resolved. Therefore, even if the perceptual enhancement driven by fine spatial attention is limited to a narrow range of lower spatial frequencies, the enhanced frequency range may shift toward higher spatial frequencies in the foveola compared to what happens extrafoveally (***Figure 1****B*). Alternatively, attention at the foveal scale might preserve its enhancement of low spatial frequencies while extending it to high spatial frequencies, leading to a broad, rather than narrow, range of modulation (***Figure 1****C*). Any of these scenarios could account for the improvement in orientation discrimination observed in ***Guzhang et al. (2021***).

**Figure 1.**
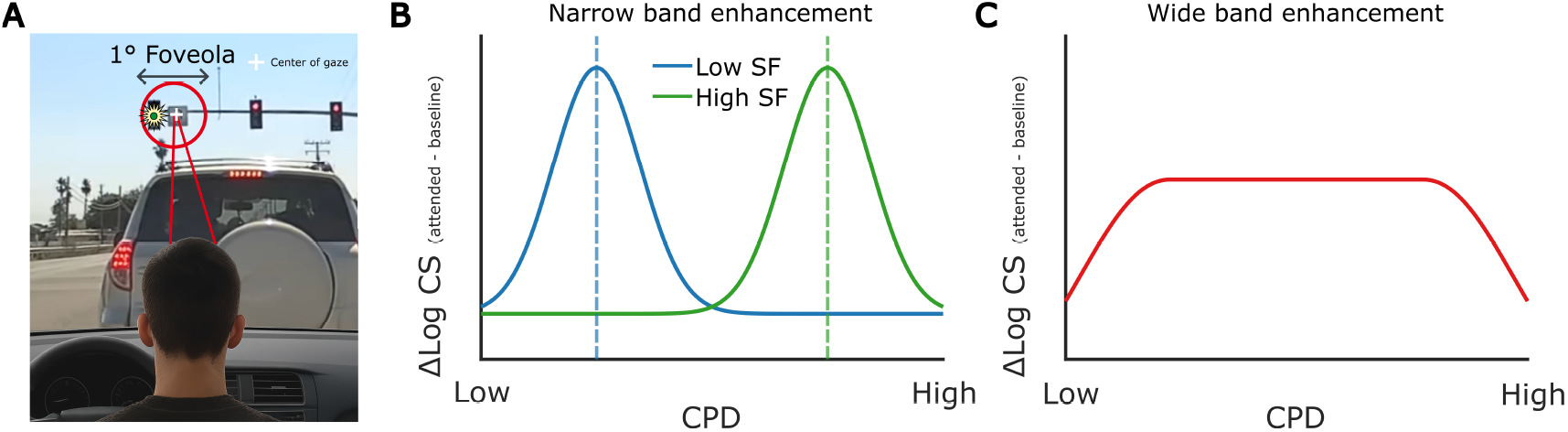
Fine-tuning exogenous attention within the foveola. (**A**) Fine-tuning of exogenous attention within the foveola occurs, for example, when we are looking at a distant traffic light—occupying less than 1° of our visual field—that suddenly turns green, capturing our attention and prompting us to move forward. As a result of the fine-tuning of exogenous attention, contrast sensitivity could be enhanced for a narrow range of spatial frequencies, centered around lower spatial frequencies (blue) or higher spatial frequencies (green) (**B**). On the other hand, contrast sensitivity may be enhanced uniformly across a wide range of spatial frequencies (**C**).

In the current study, we addressed two main questions. First, does exogenous attention at the foveolar scale enhance visual processing across a narrow or a broader range of spatial frequencies? Second, if the enhancement operated within a narrow frequency band, which range of spatial frequencies benefits the most from such fine-grained shifts of attention? Addressing these questions is crucial because, while it is now established that covert attention can be selectively shifted even within the central fovea, it remains unclear whether it follows the same *modus operandi* foveally and extrafoveally. If a similar range of spatial frequencies is enhanced by exogenous attention in both the foveola and extrafovea, it would suggest that exogenous attention operates similarly across the visual field, regardless of the spatial resolution achievable at different eccentricities. In contrast, if the modulation of spatial frequencies differs, it would indicate that the mechanisms of exogenous attention are flexibly tuned in the foveola and adjusted based on the spatial resolution that can be achieved at this scale.

Studying attentional control in the foveola presents unique challenges. Continuous microscopic eye movements during fixation cause constant displacement of the retinal input (***Martinez-Conde et al., 2004***; ***Rucci and Poletti, 2015***; ***Krauzlis et al., 2017***), making it difficult to limit visual stimulation to the desired eccentricity at this scale. This poses a significant issue when investigating covert attention in the central foveola. To address these challenges, we employed high-precision eye-tracking (***Wu et al., 2023***) combined with gaze-contingent display control (***Santini et al., 2007***) to precisely monitor gaze position throughout each trial to ensure that any effects observed are solely due to covert exogenous attention and are not driven by fixational saccades (***Hafed and Clark, 2002***; ***Yuval-Greenberg et al., 2014***; ***Shelchkova and Poletti, 2020***; ***Guzhang et al., 2024***).

## Results

To examine the effects of high-resolution exogenous attention within the fovea on visual discrimination of stimuli at different spatial frequencies, we employed a 2AFC visual discrimination task in which observers were asked to discriminate the orientation of a small Gabor patch (30^′^ x 30^′^ with an overlaying 5.4^′^ Gaussian window, tilted ±45°) 30^′^ from either left or right of the fixation marker when prompted by a response cue (***Figure 2****A*). Note that, the Gabor patches used in the current study were much smaller than those typically used in studies probing extrafoveal attention (***Rossi and Paradiso, 1995***; ***Gobell and Carrasco, 2005***; ***Herrmann et al., 2010***; ***Jigo and Carrasco, 2020***). On each trial, the orientation and phase of the Gabor patch was randomized at each location independently. Eye movements were monitored at high resolution using a digital Dual Purkinje Image eye tracker (***Wu et al., 2023***) to ensure that observers maintained the center of gaze within a 10^′^×10^′^ window around the fixation point throughout the trial (***Figure 2****D*). There were no systematic differences in fixation position between valid and neutral conditions, either horizontally (valid: 0.20^′^±0.66^′^SD; neutral: 0.21^′^±0.63^′^SD; p = 0.53) or vertically (valid: -0.52^′^±0.84^′^SD; neutral: -0.48^′^±0.82^′^SD; p = 0.34).

**Figure 2.**
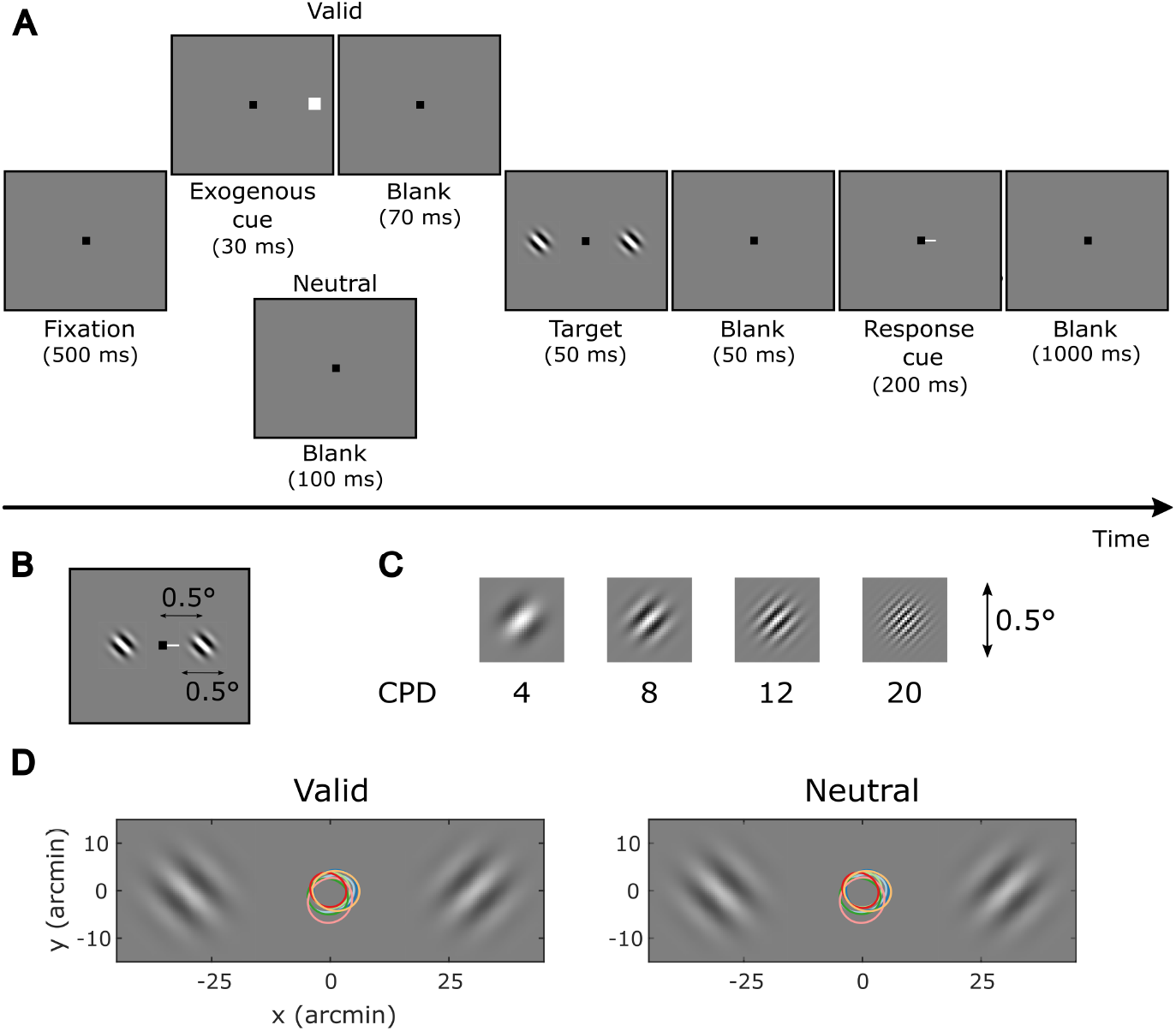
Experimental protocol. (**A**) Trials started with a fixation marker at the center of the monitor. Observers were instructed to maintain fixation at the center throughout the trial. After a brief flash of the exogenous cue to capture observers’ attention, two Gabor patches independently tilted (±45°) were briefly displayed, one on each side of the fixation marker. At the end of the trial, a response cue appeared, and observers had to report the orientation of the stimulus that was previously presented at the cued location. In valid trials, the exogenous cue and response cue indicated the same spatial location. In neutral trials, no exogenous cue was presented. Valid and neutral trials had the same probability of occurrence. (**B**) Size of the stimuli. The Gabor patches had a Gaussian window of 5.4^′^ standard deviation, creating a 30^′^ x 30^′^ visible region. (**C**) Stimuli used in the experiment. Gabor patch of all spatial frequencies tested from 4 to 20 cycles per degree (CPD). (**D**) 68% contour of the gaze probability distribution in valid and neutral conditions during Gabor presentation. Color represents individual observers. **Figure 2—figure supplement 1**. Number and proportion of trials included in the analysis after filtering. **Figure 2—figure supplement 2**. Average saccade onset relative to response cue onset.

The experiment included two cueing conditions, valid and neutral, that were randomly interleaved with equal probability within each experimental block. In the valid condition, shifts of exogenous attention were elicited by a small and brief white flash (exogenous cue) presented 100 ms before the target either on the left or the right of the visual field. The cue, with 100% validity, appeared just outside the upcoming target location. In the neutral condition, no exogenous cue was presented.

We tested four spatial frequencies (SFs), 4, 8, 12, and 20 CPDs, ***Figure 2****C*), ensuring at least one full cycle of modulation within the Gabor patch even at the lowest SF (4 CPD) (***Howell and Hess, 1978***). The highest spatial frequency (20 CPD) is close to the limit of visual resolution at the eccentricity tested here. For each spatial frequency tested, an initial threshold contrast was estimated by methods of constant stimuli, then discrimination accuracy was measured at four contrast levels around the initial threshold estimate, and one additional level at the maximum contrast to measure the asymptotic performance (see Methods). The contrast level was kept constant within each experimental block but was randomized across blocks.

To examine how high-resolution exogenous attention influences performance for stimuli at different spatial frequencies, we fitted individual psychometric curves of contrast level versus discrimination accuracy and estimated contrast thresholds for each cueing condition and spatial frequency for each observer (***Figure 3****A* shows the psychometric curves for one example observer with the result of all psychometric curves included in ***Figure 3—figure Supplement 1***).

**Figure 3.**
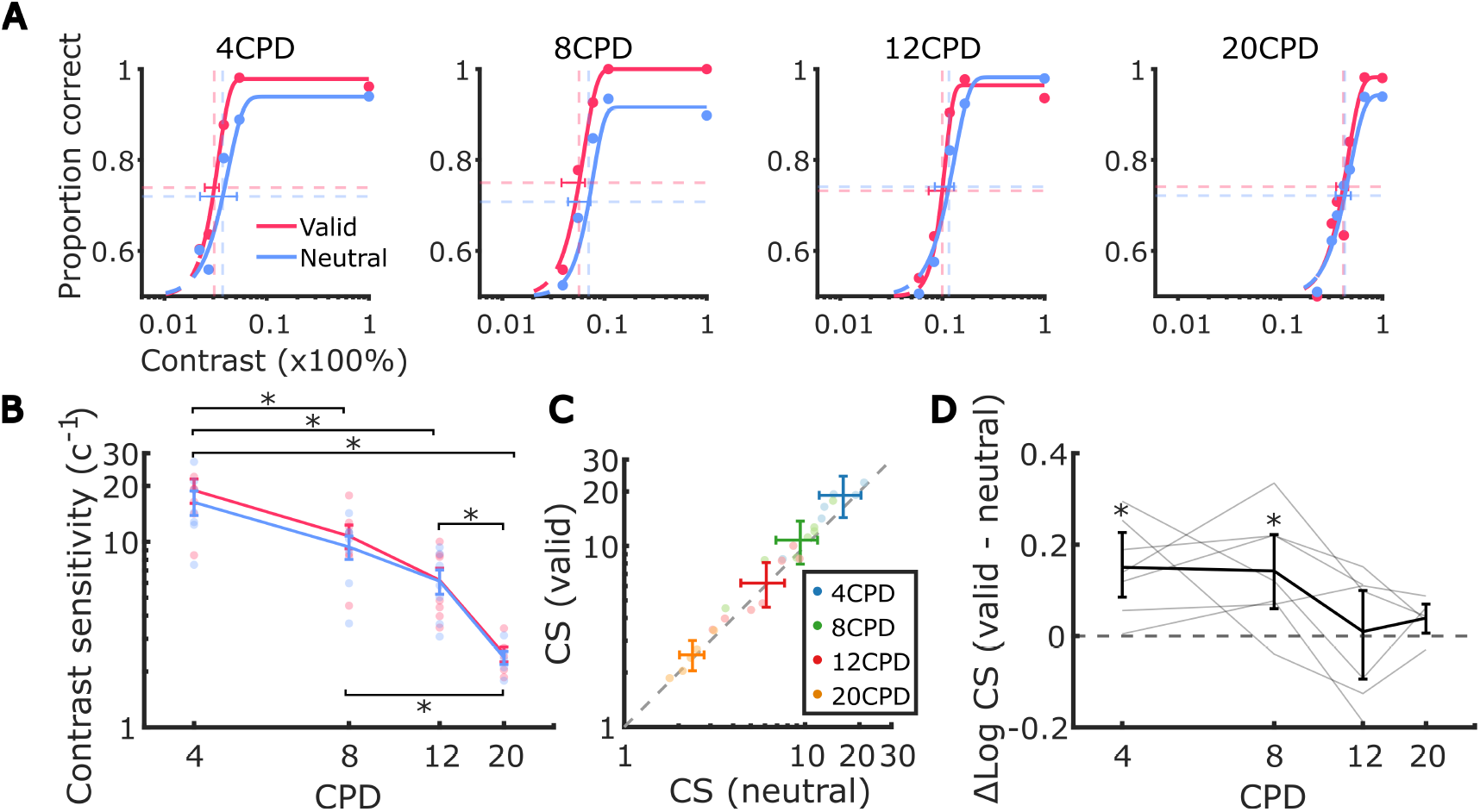
Effect of fine-grained exogenous attention on contrast sensitivity. (**A**) Psychometric functions illustrating example observers’ discrimination accuracy for Gabor patches with spatial frequencies of 4, 8, 12, and 20 cycles per degree (CPD). The size of each dot corresponds to the number of trials included at a specific contrast value. Vertical lines indicate contrast thresholds, while horizontal lines represent the accuracy level midway between chance performance and maximum performance. (**B**) Average contrast sensitivity, calculated as the inverse of the contrast threshold, across spatial frequencies in valid and neutral conditions. Each dot represents an individual observer. Error bars denote the standard error of the mean (SEM). Asterisks mark a significant difference in contrast sensitivity between pairs of spatial frequencies. (**C**) Average contrast sensitivity in neutral condition against that in valid condition at each spatial frequency. Each dot represents an individual observer. Error bars indicate the bootstrapped 95% confidence intervals. (**D**) Average difference in log-scaled contrast sensitivity between valid and neutral conditions across different spatial frequencies. Each line corresponds to the log-scaled contrast sensitivities from each observer. Error bars represent the bootstrapped confidence intervals. Asterisks mark post-hoc pairwise comparison results between valid and neutral conditions within each spatial frequency. **Figure 3—figure supplement 1**. Psychometric functions for all observers in all conditions. **Figure 3—figure supplement 2**. Contrast sensitivity results including 2 CPD. **Figure 3—figure supplement 3**. Relation between mean and variability of contrast sensitivity (CS). **Figure 3—figure supplement 4**. Attention benefit plotted as ratio of contrast sensitivity between valid and neutral trials.

We first examined the full factorial effects of spatial frequency (SF) and cueing on contrast sensitivity estimated from the observers’ psychometric functions by fitting a linear mixed-effects regression to the estimates of log-transformed contrast sensitivity at all 8 conditions (4 SF x 2 cueing) across observers. The contrast sensitivity captures the effects on the intercept and/or slope of the psychometric function.

Consistent with the literature (***Howell and Hess, 1978***; ***Bex et al., 2009***; ***Lovegrove et al., 1980***; ***Rovamo et al., 1992, 1993***; ***Pointer and Hess, 1989***), we observed a main effect of spatial frequency on contrast sensitivity (*χ*^2^(3) = 65.3, *p* < 0.0001, ***Figure 3****B*; see ***Appendix 1—Table 1*** for full model output); contrast sensitivity was highest at 4 CPD and decreased as the spatial frequency increased.^1^ We also observed a significant main effect of attention on contrast sensitivity (*χ*^2^(1) = 7.3, *p* < 0.001; mean_valid_ = 10.1 ± 7.56SD and mean_neutral_ = 9.03 ± 6.39SD): averaging across all spatial frequencies, the valid cueing condition resulted in higher contrast sensitivity (***Figure 3****B, C*). These findings indicate that exogenous attention led to a contrast gain across spatial frequencies in the attended subfoveolar region.

Notably, the improvement in contrast sensitivity driven by fine-grained attention was not uniform across spatial frequencies (***Figure 3****D*, also visible in ***Figure 3****B, C*). We observed a statistically significant interaction between spatial frequency and attention (*χ*^2^(3) = 9.3, *p* = 0.0258), indicating that contrast gains were selective. Specifically, contrast sensitivity exhibited a contrast gain in the valid condition compared to the neutral condition at lower spatial frequencies (4 and 8 CPD) (mean gain_4 CPD_ = 2.62 ± 2.13 SD, Cohen’s *d* = 0.79 and mean gain_8 CPD_ = 1.36 ± 1.25 SD, *d* = 0.75, *p*s< 0.004; see ***Appendix 1—Table 2*** for all pairwise comparisons.). However, the contrast sensitivity gains at higher spatial frequencies (12 and 20 CPD) were smaller and did not reach statistical significance (mean gain_12 CPD_ = 0.09 ± 0.91SD and mean gain_20 CPD_ = 0.11 ± 0.13SD, *p*s> 0.5). In addition to examining the contrast gain within each spatial frequency, we also compared the amount of contrast gain within each pair of spatial frequencies (see ***Appendix 1—Table 3*** for all pairwise comparisons). Post-hoc pairwise comparisons revealed that attention modulation did not differ between the two low-mid spatial frequencies (4 and 8 CPD, *p* > 0.8) or between the two mid-hight spatial frequencies (12 and 20 CPD, *p* > 0.7). Importantly, attention led to a larger enhancement of contrast sensitivity at the low-mid spatial frequencies (4 and 8 CPD) compared to 12 CPD (Δmean gain_4 CPD - 12 CPD_ = 2.53 ± 2.16SD, *d* = 0.65, and Δmean gain_8 CPD - 12 CPD_ = 1.27 ± 0.99SD, *d* = 0.62, *p*s< 0.025). When comparing the low-mid spatial frequencies (4 and 8 CPD) to the highest spatial frequency tested (20 CPD), differences in contrast gains were even larger, though these effects were not statistically significant (Δmean gain_4 CPD - 20 CPD_ = 3.35 ± 1.96SD, *p* = 0.066 and Δmean gain_8 CPD - 20 CPD_ = 1.45 ± 1.46SD, *p* = 0.084). As detailed in Methods, two observers did not have data for the 20 CPD condition. Statistical power might thus have been reduced for comparisons against this condition.

These results demonstrate that, similar to the selectivity in enhancements in visual periphery, micro-shifts of exogenous attention within the central fovea selectively enhanced contrast sensitivity primarily for low- to mid-range frequencies (4 to 8 CPD).

In addition to contrast sensitivity, we examined the effects of attention on asymptotic performance. Research on *extra*foveal vision has returned mixed results with respect to the effects of exogenous attention on asymptotic performance. It has been found that exogenous attention can enhance both contrast sensitivity (contrast gain) but also asymptotic performance, a phenomenon known as response gain (***Morrone et al., 2002, 2004***; ***Ling and Carrasco, 2006***; ***Pestilli et al., 2009***; ***Reynolds and Heeger, 2009***; ***Herrmann et al., 2010***). However, some studies have found significant effects only on contrast sensitivity, with no notable impact on asymptotic performance in extrafoveal vision (***Cameron et al., 2002***; ***Herrmann et al., 2010***; ***Jigo and Carrasco, 2020***). It has been argued that whether covert attention influences contrast gain, response gain, or both, depends on stimulus size, size of the attended area as well as spatial uncertainty(***Herrmann et al., 2010***; ***Wu et al., 2021***; ***Reynolds and Heeger, 2009***).

To examine how asymptotic performance was impacted by micro-shifts of foveal exogenous attention across spatial frequencies, we fitted a generalized linear mixed-effects regression to the estimated asymptotic performance from all 8 conditions of all observers (see ***Appendix 2—Table 1*** for full model output). Observers’ ability to discriminate the orientation at full contrast (asymptotic performance) decreased with increasing frequency (***Figure 4****B, C*), but this change was not significant (*χ*^2^(3) = 7.6, *p* = 0.0545).^2^ The main effect of attention was significant, with overall higher asymptotic performance in the valid condition compared to the neutral condition (*χ*^2^(1) = 11.9, *p* < 0.001; mean_valid_ = 0.97 ± 0.03SD and mean_neutral_ = 0.94 ± 0.04SD; see ***Figure 4****A, C*). These findings suggest that fine-grained attention resulted in a general response gain, enhancing the ability to discriminate the orientation of high-contrast stimuli.

**Figure 4.**
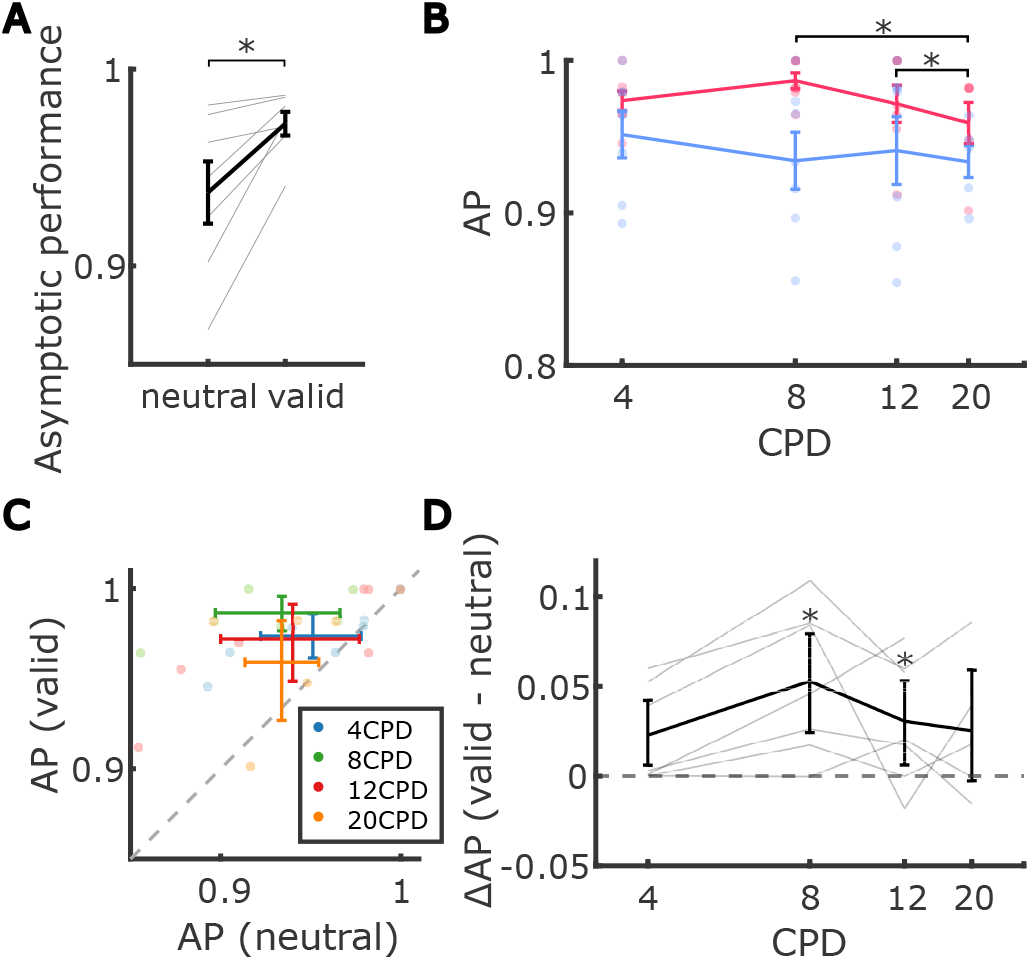
Effect of fine-grained exogenous attention on asymptotic performance. (**A**) Average asymptotic performance, defined as discrimination accuracy at maximum contrast, pooled across spatial frequencies in valid and neutral conditions. Each dot represents an individual observer. Error bars denote the standard error of the mean (SEM). (**B**) Average asymptotic performance across spatial frequencies in valid and neutral conditions. Each dot represents an individual observer. Error bars denote the standard error of the mean (SEM). Asterisks mark a significant difference in asymptotic performance between pairs of spatial frequencies. (**C**) Average asymptotic performance in the neutral condition against that in the valid condition at each spatial frequency. Each dot represents an individual observer. Error bars indicate the bootstrapped 95% confidence intervals. (**D**) Average difference in asymptotic performance between valid and neutral conditions across different spatial frequencies. Each line corresponds to an individual observer. Error bars indicate the bootstrapped confidence intervals. Asterisks mark post-hoc pairwise comparison results between valid and neutral conditions within each spatial frequency. **Figure 4—figure supplement 1**. Maximum a posteriori (MAP) estimates (points) and 95% confidence intervals for contrast sensitivity (CS) and asymptotic performance (AP).

Unlike for contrast sensitivity, we did not observe a significant interaction between spatial frequency and attention on asymptotic performance (Figs.4*D*, *χ*^2^(3) = 5.1, *p* > 0.15).^3^ Therefore, asymptotic performance exhibited a significant effect of attention alone, with no detected significant effects of spatial frequency or of the interaction between spatial frequency and attention.

## Discussion

Whereas attention is often believed to be either uniformly allocated at the center of gaze or selectively shifted to locations outside of the central fovea, recent research has shown that humans are also capable of allocating attention within the central fovea in a spatially selective manner, enhancing our ability to perceive fine spatial stimuli (***Poletti et al., 2017***; ***Guzhang et al., 2021***). Humans can focus processing resources on a specific region of the central fovea, enhancing visual processing within that small area while suppressing processing at other unattended locations just a few arcminutes away.

These findings raise the question of what spatial frequencies are enhanced by fine-grained shifts of covert attention within the foveola. In our previous work (***Guzhang et al., 2021***), observers were asked to perform a coarse orientation (±45°) discrimination task. This experiment did not manipulate spatial frequency. Subjectively, however, the stimuli in this task could be distinguished as long as frequency information of more than 3 cycles per degree (CPD) was available to the observer. The attentional gain we observed in the discrimination task could therefore have resulted from foveal exogenous attention enhancing spatial frequencies anywhere above 3 CPD. Thus, it remains unclear which spatial frequencies are enhanced when exogenous attention is allocated at the fine scale within the foveola. Additionally, it is unclear whether fine-grained attention in the foveola is governed by the same principles and is modulated similarly to extrafoveal attention, especially considering the stark differences in spatial resolution between the foveola and the rest of the visual field. The high-acuity foveola can resolve spatial frequencies up to 30 CPD, whereas just five degrees away from the center of gaze, this limit drops to around 10 CPD (***Virsu and Rovamo, 1979***). Therefore, rather than enhancing the same range of low spatial frequencies as for extrafoveal vision, fine-grained foveal attention may shift or extend its enhancement toward higher frequencies in the foveola.

Our findings indicate that fine-grained shifts of covert exogenous attention in the foveola enhance contrast sensitivity within a narrow range of spatial frequencies, peaking at low to mid frequencies (4–8 CPD). In particular, we found little or no attentional gain at higher spatial frequencies (12–20 CPD), which are closer to the limits of visual resolution at the eccentricity tested (0.3° from the preferred locus of fixation). Whereas enhancements in contrast sensitivity were relatively selective to a narrow band of spatial frequencies, overall asymptotic performance increased as a result of exogenous attention, with no detected dependence on spatial frequency. However, it is worth noting that the statistical power to detect the interaction might differ between analyses of contrast sensitivity and analyses of asymptotic performance, given that the latter tends to involve differences close to its bounds (***Bicknell et al., 2025***; ***Jaeger, 2008***).

Prior work on coarse exogenous attention shifts between central and peripheral vision has shown that peak attentional benefits for contrast sensitivity occur around 2–4 CPD, with a sizable benefit occurring also at 8 CPD, for a large foveal stimulus (***Jigo and Carrasco, 2020***). In the present study, the largest attentional gains were observed at the lowest spatial frequency tested (4 and 8 CPD). Because of the small stimulus size required to probe fine-grained attention at the foveal scale, we prioritized testing of spatial frequencies that yielded at least one full cycle within the stimulus aperture. Nevertheless, we conducted a post hoc evaluation at 2 CPD using the same Gabor size (see ***Figure 3—figure Supplement 2***). Because this stimulus contained less than one full cycle, performance may have been compromised (***Howell and Hess, 1978***). With this caveat in mind, baseline contrast sensitivity was comparable at 2 and 4 CPD and declined at 8 CPD, suggesting a plateau between 2 and 4 CPD. Attention significantly improved contrast sensitivity also at 2 CPD, with contrast gain comparable to that observed at 4 and 8 CPD (see caption of ***Figure 3—figure Supplement 2***).

Thus, the range of spatial frequencies enhanced by fine-grained, exogenously triggered attention closely mirrors that observed when attention is broadly distributed across the fovea. The magnitude of the contrast gain (≈ 20%) was likewise consistent with prior findings (***Jigo and Carrasco, 2020***) and slightly larger than that reported by (***Herrmann et al., 2010***) (see ***Figure 3—figure Supplement 4***).

### Contrast vs. response gain

Previous studies examining attentional modulation of contrast response functions in extrafoveal vision have reported heterogeneous gain profiles, with some observing predominantly contrast gain (e.g., ***Cameron et al., 2002***; ***Jigo and Carrasco, 2020***) and others reporting a mixture of contrast and response gain (e.g., ***Ling and Carrasco, 2006***; ***Pestilli et al., 2009***). This variability has been attributed to differences in stimulus size and the spatial extent of the attentional field (***Reynolds and Heeger, 2009***; ***Herrmann et al., 2010***). Within the normalization model of attention, attention multiplicatively scales stimulus drive before normalization and therefore affects both the stimulus drive and, indirectly, the pooled suppressive drive. When the attentional field is small relative to the stimulus, the attended stimulus drive is amplified more than the suppressive drive, producing a multiplicative upward scaling of the contrast response function, with the largest effects at high contrast. On the other hand, when the attentional field is large relative to the stimulus, stimulus and suppressive drive are modulated more proportionally, shifting the effect toward contrast gain. It has also been shown that the effects of endogenous attention manifest as contrast gain, and the effects of exogenous attention can manifest as a mixture between contrast and response gain (***Ling and Carrasco, 2006***; ***Pestilli et al., 2009***). Hence, the detection of both response and contrast gain in our paradigm, although different from ***Jigo and Carrasco (2020***), is in line with this expectation.

Further, foveal neurons have smaller integration fields and reduced spatial pooling than peripheral neurons. At the retinal level, classic work on macaque midget ganglion cells showed that receptive fields in the central fovea are extremely small and can be dominated by single-cone input, consistent with minimal spatial pooling at the very center of gaze (***Dacey and Petersen, 1992***; ***Dacey, 1993***). This characterization seems to be maintained at the LGN (***Ramsey et al., 2026***). More generally, human EEG studies using steady-state visual evoked potentials, as well as behavioral studies, have shown that surround suppression is stronger in the visual periphery than near the fovea (***Xing and Heeger, 2000***; ***Petrov et al., 2005***; ***Vanegas et al., 2015***). Although there has been relatively little direct electrophysiological work systematically comparing suppressive zone size or normalization strength between central fovea and peripheral neurons in primate visual cortex, the available evidence raises the possibility that the normalization pool engaged at the foveal scale is smaller and less influential than in extrafoveal vision. In the normalization model of attention the resulting attention gain profile depends on the balance between the attended stimulus drive and the pooled suppressive drive (***Reynolds and Heeger, 2009***). If suppressive interactions are indeed weaker within the foveola, then directing attention to a specific foveal locus may amplify the local stimulus drive more than the pooled suppressive drive, thereby favoring response-gain-like effects over contrast-gain-like effects.

Under this interpretation, the detection of response gain in our data may reflect the unusually small integration and suppressive fields engaged by fine-scale attention within the foveola, in contrast with the seemingly more pronounced contrast-gain effects reported in ***Jigo and Carrasco (2020***). This possibility may also help explain why contrast-gain-like effects were less pronounced at higher spatial frequencies in our study, since high spatial frequency stimuli are expected to engage smaller receptive fields and even less spatial integration, further reducing the contribution of normalization (***Teichert et al., 2007***; ***Serrano-Pedraza et al., 2012***). This account—based on posthoc between-study comparisons—remains speculative. Future work will be needed to determine whether fine-scale attention within the fovea is mediated by mechanisms distinct from those operating when attention is distributed more broadly across the central fovea, and whether such differences alter the size or influence of the normalization pool.

### Alternative explanations for the observed effects

While our primary goal was to examine spatially localized, exogenously triggered fine-grained covert attention within the foveola, one could argue that temporal cueing may have contributed to the observed effects. Specifically, even though the timing of the stimulus onset was fixed across trials, the shorter interval between the additional exogenous cue and the target in valid trials could have acted as a temporal warning signal, potentially enhancing performance relative to neutral trials (***Duyar et al., 2023***). While we cannot entirely rule out temporal cueing effects, their influence is likely limited. Because stimulus timing was fixed and observers completed many trials, the temporal contingency between the cue and stimulus was likely overlearned. Thus, although a minor temporal benefit may have been present in valid trials, the primary attentional advantage observed here is most attributed to spatially localized attentional engagement at the cued location.

Besides the possible contribution of temporal attention, endogenous attention may also have influenced the results. Although our cue was salient and abrupt in onset, and therefore likely triggered an involuntary exogenous shift of attention, its 100% validity could in principle have supported endogenous maintenance following the initial orienting response (***Chica et al., 2013***). However, endogenous attentional benefits typically emerge on a slower timescale, with onset latencies of approximately 200 ms or longer (***Carrasco, 2011***). In contrast, the cue–target SOA in our study was only 100 ms, and the target itself was presented for 50 ms. Under these temporal constraints, any endogenous contribution would be expected to arise only after target offset. Thus, the attentional benefits reported here likely reflect rapid, stimulus-driven exogenous mechanisms.

### Future directions

In addition to its perceptual consequences, fine-grained exogenous attention within the foveola may play a preparatory role in oculomotor behavior. When examining the trials that were excluded from main analyses, in which observers happened to perform a saccade following the response cue, we observed a significant reduction in saccade onset latency in valid compared to neutral trials (***Figure 2—figure Supplement 2***). This effect suggests that exogenous attentional deployment within the foveola not only enhances visual sensitivity but may also facilitate the rapid initiation of gaze shifts toward salient events. Although the present task was deliberately constrained, requiring strict fixation during stimulus presentation, this finding hints at a functional coupling between fine-grained exogenous attention within the foveola and the preparation of subsequent eye movements. Future studies using more naturalistic viewing conditions, in which observers are free to move their eyes, will be critical for determining whether this preparatory mechanism serves to efficiently guide microsaccades or saccades toward behaviorally relevant stimuli in everyday vision.

In everyday life, covert exogenous attention is often engaged when a salient stimulus captures our focus. This evolutionarily important mechanism ensures that we continuously monitor our environment for unexpected events and prepare to respond accordingly (***Yantis, 1993***; ***Theeuwes, 2010***). Our previous work has demonstrated that attentional shifts can also occur locally within the high-acuity foveola. Here, we show that these fine-grained attention shifts function similarly to those in the extrafoveal region, enhancing visual sensitivity to coarse stimulus features. This mechanism is essential for everyday tasks, such as driving or reading. Our findings not only shed light on the functionality of fine-grained covert attention within the foveola but also reinforce the idea that exogenous attention operates under similar principles as extrafoveal vision. Specifically, exogenous attention remains an inflexible mechanism for selective processing—even in the foveola, where higher spatial frequency information is available, it does not enhance contrast sensitivity of the finer details but instead prioritizes coarser stimulus features. Functionally, this selective enhancement of contrast sensitivity at low to mid spatial frequencies provides a preview of small but salient stimuli located just a few arcminutes from the preferred locus of fixation in everyday tasks. By enhancing contrast sensitivity of these stimuli before direct fixation, this mechanism enables the visual system to rapidly assess their relevance and guide the planning of microsaccades, ensuring efficient and precise shifts of gaze to bring these stimuli into the foveal region for detailed examination.

## Methods and Materials

### Observers

7 human observers in total, 6 emmetropic observers, and 1 observer with 20/20 corrected vision participated in the experiments (4 females, 3 males; age range 18 - 27 years old). The experiment was approved by the University of Rochester Institutional Review Boards. The experimenter reviewed and explained the material in the consent form to the observers before conducting the experiment. The form was signed only after the participant fully understood the material and voluntarily agreed to take part in the study. Consent was obtained from all observers in the study.

### Stimuli and Apparatus

Stimuli were displayed on an LCD monitor (ASUS ROG SWIFT 360Hz PG259QN) at a refresh rate of 360 Hz and spatial resolution of 1920 x 1080 pixels. Observers performed the task monocularly with their right eye while the left eye was patched. A dental-imprint bite bar and a headrest were used to prevent head movements. Eye movements were recorded with high precision using a custom-made digital Dual Purkinje Image (dDPI) eye tracker, which has a sampling rate of 1 kHz (***Wu et al., 2023***). The system has an internal noise well below 1^′^ and a spatial resolution of 1^′^ (***Ko et al., 2016***; ***Wu et al., 2023***). Stimuli were rendered using EyeRIS, a custom-developed system that allows flexible gaze-contingent display control (***Santini et al., 2007***). This system acquires eye movement signals from the eye tracker, processes them in real time, and updates the stimulus on the display according to the desired combination of estimated oculomotor variables.

### Procedure and Experimental Task

#### Calibration

Every session started with the setup of the bite bar. A magnetized helmet was used to position the observer’s head. When accurate localization of gaze position is necessary, calibration represents an important stage of the experimental procedure, which was performed in two phases, automatic calibration followed by manual calibration. During automatic calibration, observers sequentially fixated on each of the nine points of a 3-by-3 grid, as is customary in all oculomotor experiments. After completing automatic calibration, observers were instructed to perform a manual calibration where they refined the pixel-to-pixel mapping, given by the automatic calibration. To this end, observers fixated again on each of the nine points of the grid while the location of the line of sight was displayed in real time on the screen. Observers used a joypad to correct the predicted gaze location, shifting the real-time display to align with the grid point for each fixation, if necessary. These corrections were then incorporated into the transformation of the gaze position as well. This dual-step calibration procedure allows more accurate localization of gaze position than standard single-step procedures. A similar manual calibration procedure was repeated before each trial but only for the central fixation location to compensate for unpreventable head movements.

#### Experimental task

Observers were instructed to fixate on a central marker (5 by 5 arcminutes) throughout each trial. On valid trials, an exogenous cue—a white square (8-by-8 arcminutes)—appeared 500ms after fixation. The cue appeared for 30 ms at 0.75 deg eccentricity to the left/right of the fixation marker, with each location occurring randomly with equal probability. The smaller exogenous cue, positioned offset from the Gabor patch, was used to prevent forward masking and ensure clear perception of the Gabor patch. Shortly after the cue disappeared (70 ms), two small Gabor patches (0.5 deg visible area), tilted +/- 45 degrees, with a phase of 0 or 90 degrees were shown (50 ms) on the left/right side at 0.5 deg eccentricity. The tilt of the two Gabor patches was randomly and independently chosen on each trial. And the phase was randomly selected on each trial but consistent between the two patches. The spatial frequency of the Gabors was 4, 8, 12, or 20 CPD. After the stimulus offset, a response cue was presented, and observers were instructed to report the orientation of the stimulus previously presented at that location. The trial concluded either when observers responded or automatically after 1000 ms if no response was given following the appearance of the response cue. On valid trials, the response cue always indicated the same location as the exogenous cue, making the cue 100% valid. On neutral trials, no exogenous cue was presented, and the response cue indicated one of the two possible locations randomly.

When observers first arrived for the study, they were given task instructions and completed 50 familiarization trials to become accustomed to the setup and the task. Following this, because contrast sensitivity varies considerably across SF and eccentricity (***Rovamo et al., 1992***), for each spatial frequency tested, observers underwent a preliminary session in which an initial estimate of contrast threshold, defined as the contrast needed to achieve 70% discrimination accuracy in the neutral condition. It typically took around 50-100 trials to find the target contrast threshold for each spatial frequency. Neither the familiarization trials nor the threshold-estimation trials were included in the final analyses. After the thresholds were obtained, each observer was tested at five different contrast values around the estimated threshold. One of these values included presenting the grating at 100% contrast to obtain a precise estimate for the upper-performance asymptote. The remaining four levels were ±0.075 and ±0.225 log_10_ units from the initial threshold estimate.

If the initial estimates were within 0.225 log_10_ units of 100% contrast (i.e., >= 60% contrast), the rest of the four contrast values were -0.6, -0.45, -0.3, and -0.15 log_10_ units compared to the initial estimate (***Prins, 2012***). Within each experimental session, a single spatial frequency was tested, and the corresponding contrast levels were presented in a block design. All five contrast levels for a given spatial frequency were tested within a single experimental session. Observers completed 100 trials per contrast level. And each spatial frequency was tested twice on two separate days. Therefore, observers completed approximately 4000 trials in total. The order of spatial frequency tested was randomized across observers.

Two observers were not tested at 20 CPD because their performance remained at chance level even with gratings at maximal contrast. It is possible given that 20 CPD was near the visual resolution limit at the tested eccentricity.

### Data Analysis

#### Eye movements

Only trials with uninterrupted tracking in which the fourth Purkinje image was never eclipsed by the pupil margin, were selected for data analysis. Trials in which the gaze was > 10’ away from the center position 50 ms before the onset of the exogenous cue *t*_0_ 50 ms after the offset of the target, and trials with blinks, saccades, or microsaccades occurring at any time during the period of interest (50 ms before the onset of the exogenous cue to 200 ms after the offset of the Gabor patches), were discarded. Periods of blinks were automatically detected by the dDPI eye tracker. Eye movements with a minimal amplitude of 30’ and a peak velocity higher than 3°/s were categorized as saccades. Saccades with an amplitude of less than 0.5° (30’) were defined as microsaccades. Saccade amplitude was defined as the vector connecting the point where the speed of the gaze shift grew greater than 3°/s (saccade onset) and the point where it became less than 3°/s (saccade offset). Periods that were not classified as saccades or blinks were labeled as drifts. Observers had 1000 ms to respond, and trials were excluded from further analysis if observers responded too fast (< 100 ms) or too slow (> 1000 ms), resulting in the exclusion of 0.02% ± 0.02% of the trials. Approximately between 50 and 100 trials per contrast level per cueing condition were retained after filtering. ***Table 1*** summarizes the data remaining for analysis for each condition (see ***Figure 2—figure Supplement 1*** for a detailed breakdown by observer).

**Table 1.** Average number and percentage of trials retained for analysis after filtering (mean ± SE across observers) across different spatial frequencies and cueing conditions. Brackets indicate bootstrapped 95% confidence intervals. Observers who were unable to perform the task at 20 CPD were excluded from the 20 CPD trial counts.

| SF (CPD) | Valid |  | Neutral |  |
| --- | --- | --- | --- | --- |
|  | Count | % | Count | % |
| 4 | 514 [474, 549] | 79.1 [66.1, 90.4] | 518 [486, 548] | 79.6 [67.3, 90.1] |
| 8 | 491 [417, 551] | 82.0 [73.4, 89.3] | 489 [418, 553] | 81.8 [73.3, 89.6] |
| 12 | 487 [416, 552] | 80.7 [72.9, 88.0] | 507 [444, 567] | 84.1 [77.5, 90.6] |
| 20 | 531 [504, 556] | 86.8 [82.9, 90.9] | 528 [497, 554] | 86.4 [81.1, 91.6] |

#### Psychometric function fitting

Weibull functions were fitted to the responses of the orientation discrimination task, using the maximum likelihood procedure implemented in the psignifit 4 toolbox (***Schütt et al., 2016***) for MATLAB. Separate functions were fitted for each combination of observer, attention conditions (attended and neutral), and spatial frequency, for a total of 8 psychometric function fits (2 attention conditions x 4 spatial frequencies) per observer. Each fit resulted in maximum a posteriori (MAP) estimates for the intercept *α*, slope *β*, threshold *θ*, and lapse rate *λ* of the psychometric function (the guess rate *γ* was set to .5, given the 2AFC task). Two estimates were extracted from the MAP estimates to examine the effects of fine-grained exogenous attention across spatial frequencies — contrast sensitivity and asymptotic performance. Contrast sensitivity was defined as the inverse of the threshold (the midpoint on the psychometric curve between chance performance and maximum performance). Asymptotic performance was calculated by subtracting the lapse rate from 1, representing the discrimination accuracy at the highest contrast level of the stimuli. In total, this procedure resulted in 52 estimates each of contrast sensitivity and asymptotic performance (5 observers with 4 SFs x 2 cueing conditions, and 2 observers, who did not complete the 20 CPD condition, with 3 SFs x 2 cueing conditions).

#### Statistical testing

Contrast sensitivity (CS) and asymptotic performance (AP) were both analyzed with (different types of) mixed-effects regressions. Each of the mixed-effects regressions contained cueing, spatial frequency, and their interactions as fixed-effects predictors. Cueing was effect-coded (“attended” = .5 vs. “neutral” = -.5), and frequency was coded using sliding difference, comparing the effects for each spatial frequency against the next highest spatial frequency (4 vs. 8, 8 vs. 12, 12 vs. 20). Following the recommended procedure (***Lohse, 2022***), we included the maximal possible random effect structure: random intercepts by observer, by unique combination of observer and cueing condition, and by unique combination of observer and spatial frequency condition.

CS is a bounded variable with a natural limit in that it cannot be lower than zero. Importantly, the variance of bounded variables tends to systematically decrease as their mean approaches the bound. This violates the assumption of homoskedasticity—the idea that variance should be independent of the mean and thus remain roughly constant across different conditions — an assumption that is shared by widely used statistical methods like *t*-tests, ANOVA (analysis of variance) and linear mixed-effects models (LMMs). When this assumption is violated, it can impact the reliability of statistical conclusions, affecting both Type I errors (false positives) and Type II errors (false negatives) (***Jaeger, 2008***).

Indeed, we observed a strong positive correlation between the mean and variance of contrast sensitivity: smaller variances for smaller means (***Figure 3—figure Supplement 3***). To address this issue, we log-transformed CS before analyzing it with an LMM using the lmer function from the lme4 package (***Bates et al., 2025***) in R (***R Core Team, 2024***). This largely mitigated the heteroskedasticity, except potentially in the 20 CPD condition (see ***Figure 3—figure Supplement 3***). As a precautionary measure, we verified that all main findings remained unchanged when this condition was excluded. This included the critical interaction between spatial frequency and attention (*χ*^2^(2) = 6.6, *p* < .04), which remained statistically significant.

AP is bounded both at the lower and the upper end (as it cannot be larger than 1, or smaller than the guess rate). Following recommended procedure, we thus normalized asymptotic performance to the range between 0 and 1, and analyzed it with a mixed-effects Beta model (with a logit link) using the `glmmTMB` function of the package `glmmTMB` (***McGillycuddy et al., 2025***).

Post-hoc pairwise comparisons for both mixed-effects analyses were conducted by estimating the relevant marginal means of the fitted mixed-effects regression, using the `emmeans` package (***Lenth et al., 2025***) in R. Cohen’s d (***Cohen, 1988***) of the pairwise differences was computed using the function `t_to_d` from the package `effectsize` (***Ben-Shachar et al., 2020***) in R.

#### Power analysis

Our initial sample size estimate was an approximation based on our previous study using a similar design to examine exogenous attention during high-acuity stimulus discrimination within the foveola (***Guzhang et al., 2021***). To formalize this estimate, we conducted a post-hoc power analysis following approaches used in prior work on *extra*foveal attention (***Jigo and Carrasco, 2020***) to estimate the sample size required to detect attentional effects on contrast gain. We assumed effect sizes comparable to those observed in our previous study (***Guzhang et al., 2021***). Specifically, we followed a bootstrap approach (***McConnell and Vera-Hernández, 2015***; ***Jigo and Carrasco, 2020***) in which data from two to twelve observers were randomly sampled with replacement from ***Guzhang et al. (2021***). For each resampled dataset, we conducted a one-way repeated-measures analysis of variance (ANOVA) with attention as a within-subject factor. This procedure was repeated 10,000 times, and a distribution of p-values was constructed for the main effect of attention. Statistical power was estimated as the proportion of iterations yielding a significant main effect (p < 0.05) for each sample size. The results indicated that a sample size of five observers was sufficient to achieve statistical power greater than 80% for detecting the main effect of attention. Based on this simulation, the sample size of seven observers used in the present study should provide adequate statistical power.

We conducted additional post-hoc power analyses to evaluate the power of our design to detect main effects and their interactions, using the *simr* function of the package *simr* in R. We estimated statistical power for mixed-effects models through model-based simulation by generating synthetic datasets based on the fixed and random effects structure of the fitted model, preserving the observed effect sizes and variance components. For each simulated dataset, the model was refitted, and the effect of interest was tested. By repeating this procedure 501 times across different sample sizes, power was estimated as the proportion of simulations in which the effect was statistically significant (p<0.05). Consistent with the bootstrap power analysis reported above, the results show that our study had high power (>95%) to detect the main effects of attention and spatial frequency, and moderate power (>65%) to detect the interaction. Because classical post hoc power analyses tend to be circular and anti-conservative (***Quach et al., 2022***; ***Heinsberg and Weeks, 2022***), we also used the same approach while allowing for potential overestimation of effect sizes and underestimation of variance components. Under deliberately conservative assumptions, specifically, effect sizes reduced to 75% of those observed and variance components increased by 25%, power to detect the main effects remained above 85%, while power to detect the interaction (which was nevertheless observed) was approximately 50%.

Below, we summarize the key considerations for interpreting our results.

### Limitations

Here, we analyzed log-transformed CS and AP in two separate analyses, each conducted over the observer-level estimates for each condition. We did so both (1) because this approach remains the standard in psychophysics—including in research on the role of covert attention (***Cameron et al., 2002***; ***Pestilli and Carrasco, 2005***; ***Herrmann et al., 2010***; ***Li et al., 2021***)—and (2) because the alternative would have required fitting mixed-effects *trial-level* psychometric models to the combined data from all conditions and observers (an approach that is computationally demanding, and has not yet been broadly validated). The approach taken here and in prior work on extrafoveal attention does, however, have several known limitations, some of which might be of particular relevance to questions about the effects of attention. We summarize these potential downsides here, so that they can be considered in the interpretation of our results, and addressed in future work.

First, reducing each observer’s performance in a given condition to the best-fitting psychometric parameters (*e.g*., sensitivity or asymptotic performance) ignores the uncertainty associated with those estimates. Subsequent repeated-measures ANOVAs or, in our case, mixed-effects regression analyses performed on these point estimates cannot take advantage of the information present in the trial-level data. As a result, two parameter estimates with very different levels of uncertainty (see ***Figure 4—figure Supplement 1***) may have similar influence on the analysis despite substantial differences in precision. Importantly, the precision of psychometric estimates can vary across observers and conditions because the amount and distribution of usable data differ, and because the sampled stimuli may probe different portions of the psychometric function. Ultimately, failure to adequately account for uncertainty can lead to miscalibrated inference, increasing the risk of both Type I and Type II errors.

Second, the standard approach ignores the interdependence among psychometric parameters. In practice, the same data can often be fit nearly equally well by different combinations of parameter values, for example, a steeper slope paired with a lower lapse rate, or a shallower slope paired with a higher asymptote. In particular, when performance fails to adequately constrain the upper asymptote, parameters such as threshold, slope, and lapse or asymptotic performance can become difficult to disentangle. When only the best-fitting parameter values are carried forward, all information about such covariance between parameters is lost. As a result, apparent differences in whether attention affects contrast gain or response gain may partly reflect parameter tradeoffs rather than genuine differences across studies.

Future research could employ alternative analysis approaches. In particular, it is now possible to fit mixed-effects psychometric models to the trial-level data from all conditions and all observers (***Prins, 2024***; ***Tan and Jaeger, 2025***). While this approach is computationally more demanding and requires familiarity with nonlinear mixed-effects modeling, it allows statistical tests that avoid the downsides described above.

## Data Availability

The dataset and analysis code are available on Open Science Framework (OSF): https://osf.io/pg7hd/overview?view_only=48996efa1f7641b6863b9d6d333ec31d

## Additional Information

### Author Contributions

Y.G. and M.P. designed research; Y.G. performed research; Y.G. and T.J. analyzed data; all authors drafted the paper and approved the final version of the manuscript.

## Acknowledgment

The authors wish to thank Nora Rooney for assisting with data collection.

## Appendix 1

### Effects of attention and spatial frequency on contrast sensitivity

**Appendix 1—table 1.** Linear mixed-effects model predicting log-transformed contrast sensitivity from attention, spatial frequency, and their interaction.

| Fixed Effect | Estimate | SE | 95% CI | df | t | p |
| --- | --- | --- | --- | --- | --- | --- |
| Intercept | 1.879 | 0.148 | [1.583, 2.175] | 5.96 | 12.73 | <0.001 |
| Attention (valid vs. neutral) | 0.084 | 0.028 | [0.029, 0.139] | 5.54 | 2.99 | 0.027 |
| Frequency (8 vs. 4) | -0.566 | 0.083 | [-0.729, -0.403] | 15.95 | -6.85 | <0.001 |
| Frequency (12 vs. 8) | -0.485 | 0.083 | [-0.648, -0.322] | 15.95 | -5.87 | <0.001 |
| Frequency (20 vs. 12) | -0.997 | 0.093 | [-1.179, -0.815] | 16.04 | -10.73 | <0.001 |
| Attention × Frequency (8 vs. 4) | -0.008 | 0.054 | [-0.114, 0.098] | 15.53 | -0.14 | 0.889 |
| Attention × Frequency (12 vs. 8) | -0.134 | 0.054 | [-0.240, -0.028] | 15.53 | -2.48 | 0.025 |
| Attention × Frequency (20 vs. 12) | 0.022 | 0.060 | [-0.096, 0.140] | 16.46 | 0.37 | 0.714 |

**Appendix 1—table 2.** Simple effects of attention on contrast sensitivity at each spatial frequency (SF) and their effect sizes in Cohen’s d.

| SF | Estimate | SE | 95% CI | df | t | p | Cohen's $d$ | Estimate (ratio) |
| --- | --- | --- | --- | --- | --- | --- | --- | --- |
| 4 | 0.151 | 0.043 | [0.061, 0.240] | 19.76 | 3.52 | 0.002 | 0.79 | 1.16 |
| 8 | 0.143 | 0.043 | [0.054, 0.233] | 19.76 | 3.34 | 0.003 | 0.75 | 1.15 |
| 12 | 0.009 | 0.043 | [-0.080, 0.099] | 19.76 | 0.22 | 0.830 | 0.05 | 1.01 |
| 20 | 0.032 | 0.051 | [-0.074, 0.137] | 21.37 | 0.62 | 0.540 | 0.13 | 1.03 |

**Appendix 1—table 3.** Interaction contrasts comparing attention effects between spatial frequency (SF) pairs and their effect sizes in Cohen’s d.

| SF Pair | Estimate | SE | 95% CI | df | t | p | Cohen's $d$ |
| --- | --- | --- | --- | --- | --- | --- | --- |
| 4 - 8 | 0.008 | 0.054 | [-0.107, 0.122] | 16.0 | 0.14 | 0.889 | 0.04 |
| 4 - 12 | 0.141 | 0.054 | [0.027, 0.256] | 16.0 | 2.62 | 0.019 | 0.65 |
| 4 - 20 | 0.119 | 0.061 | [-0.009, 0.247] | 16.9 | 1.97 | 0.066 | 0.48 |
| 8 - 12 | 0.134 | 0.054 | [0.019, 0.248] | 16.0 | 2.48 | 0.025 | 0.62 |
| 8 - 20 | 0.111 | 0.061 | [-0.016, 0.239] | 16.9 | 1.84 | 0.084 | 0.45 |
| 12 - 20 | -0.022 | 0.061 | [-0.150, 0.106] | 16.9 | -0.37 | 0.717 | -0.09 |

## Appendix 2

### Effects of attention and spatial frequency on asymptotic performance

**Appendix 2—table 1.** Generalized linear mixed-effects model predicting normalized asymptotic performance from attention, spatial frequency, and their interaction.

| Fixed Effect | Estimate | SE | 95% CI | z | p |
| --- | --- | --- | --- | --- | --- |
| Intercept | 2.676 | 0.349 | [1.99, 3.36] | 7.67 | <0.001 |
| Attention (valid vs. neutral) | 0.960 | 0.178 | [0.61, 1.31] | 5.41 | <0.001 |
| Frequency (8 vs. 4) | 0.289 | 0.353 | [-0.40, 0.98] | 0.82 | 0.414 |
| Frequency (12 vs. 8) | -0.181 | 0.351 | [-0.87, 0.51] | -0.52 | 0.606 |
| Frequency (20 vs. 12) | -1.020 | 0.394 | [-1.79, -0.25] | -2.59 | 0.010 |
| Attention × Frequency (8 vs. 4) | 1.127 | 0.499 | [0.15, 2.11] | 2.26 | 0.024 |
| Attention × Frequency (12 vs. 8) | -0.820 | 0.506 | [-1.81, 0.17] | -1.62 | 0.105 |
| Attention × Frequency (20 vs. 12) | -0.167 | 0.493 | [-1.13, 0.80] | -0.34 | 0.734 |

**Appendix 2—table 2.**
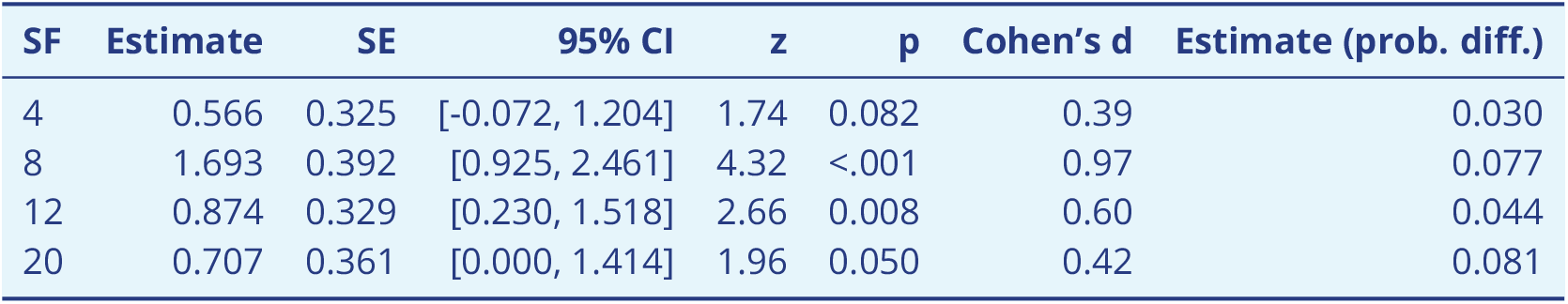
Simple effects of attention on asymptotic performance at each spatial frequency (SF) and their effect sizes in Cohen’s d.

| SF | Estimate | SE | 95% CI | z | p | Cohen's $d$ | Estimate (prob. diff.) |
| --- | --- | --- | --- | --- | --- | --- | --- |
| 4 | 0.566 | 0.325 | [-0.072, 1.204] | 1.74 | 0.082 | 0.39 | 0.030 |
| 8 | 1.693 | 0.392 | [0.925, 2.461] | 4.32 | <.001 | 0.97 | 0.077 |
| 12 | 0.874 | 0.329 | [0.230, 1.518] | 2.66 | 0.008 | 0.60 | 0.044 |
| 20 | 0.707 | 0.361 | [0.000, 1.414] | 1.96 | 0.050 | 0.42 | 0.081 |

**Appendix 2—table 3.** Interaction contrasts comparing attention effects between spatial frequency (SF) pairs on asymptotic performance and their effect sizes in Cohen’s d.

| SF Pair | Estimate | SE | 95% CI | z | p | Cohen's $d$ |
| --- | --- | --- | --- | --- | --- | --- |
| 4 - 8 | -1.127 | 0.499 | [-2.106, -0.149] | -2.26 | 0.024 | -0.56 |
| 4 - 12 | -0.308 | 0.460 | [-1.209, 0.593] | -0.67 | 0.503 | -0.17 |
| 4 - 20 | -0.140 | 0.488 | [-1.098, 0.817] | -0.29 | 0.774 | -0.07 |
| 8 - 12 | 0.820 | 0.506 | [-0.173, 1.812] | 1.62 | 0.105 | 0.40 |
| 8 - 20 | 0.987 | 0.537 | [-0.066, 2.040] | 1.84 | 0.066 | 0.45 |
| 12 - 20 | 0.167 | 0.493 | [-0.798, 1.133] | 0.34 | 0.734 | 0.08 |

**Figure 2—figure supplement 1.**
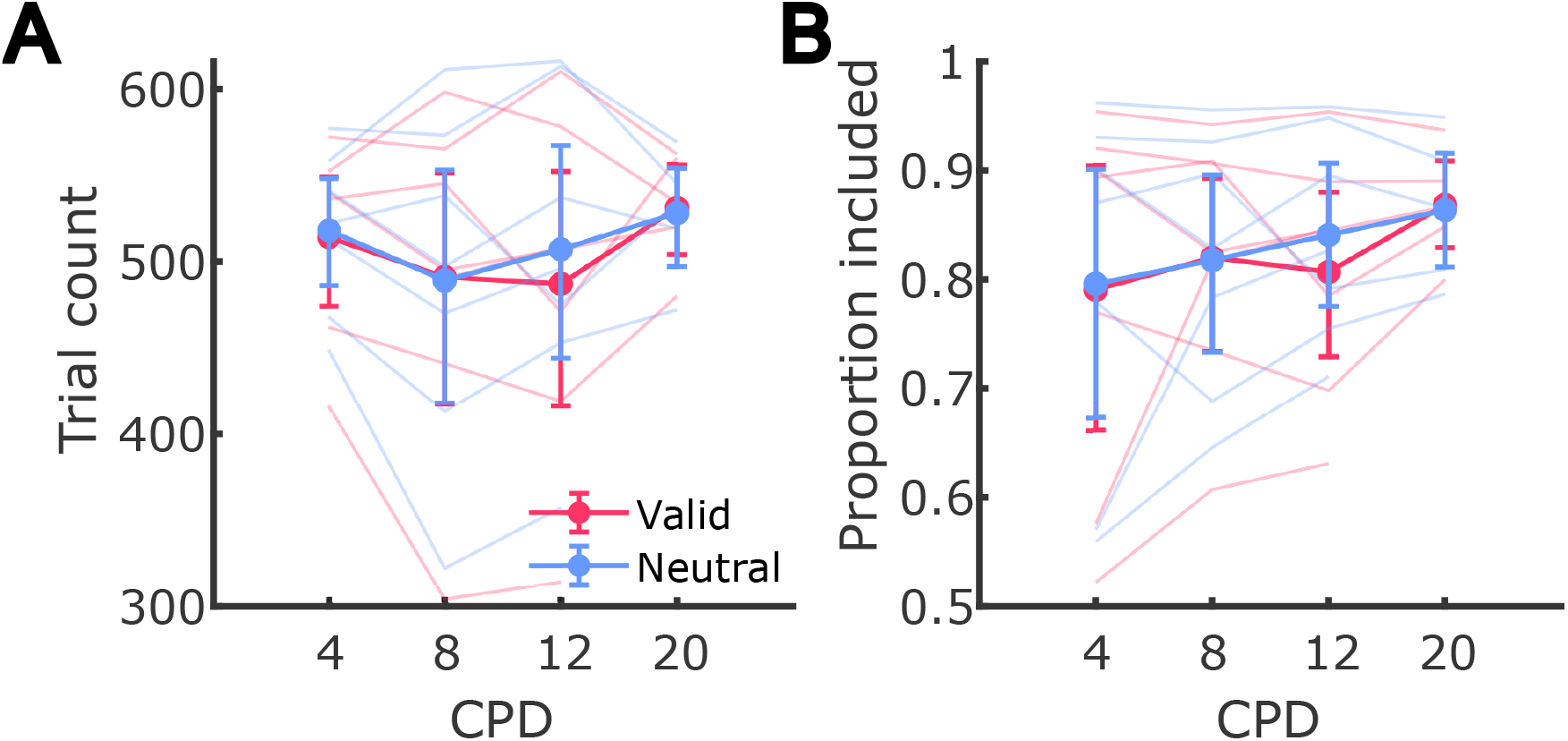
Number of trials included for analysis by condition. Each line is an observer. Error bars show the bootstrapped 95% confidence intervals.

**Figure 2—figure supplement 2.**
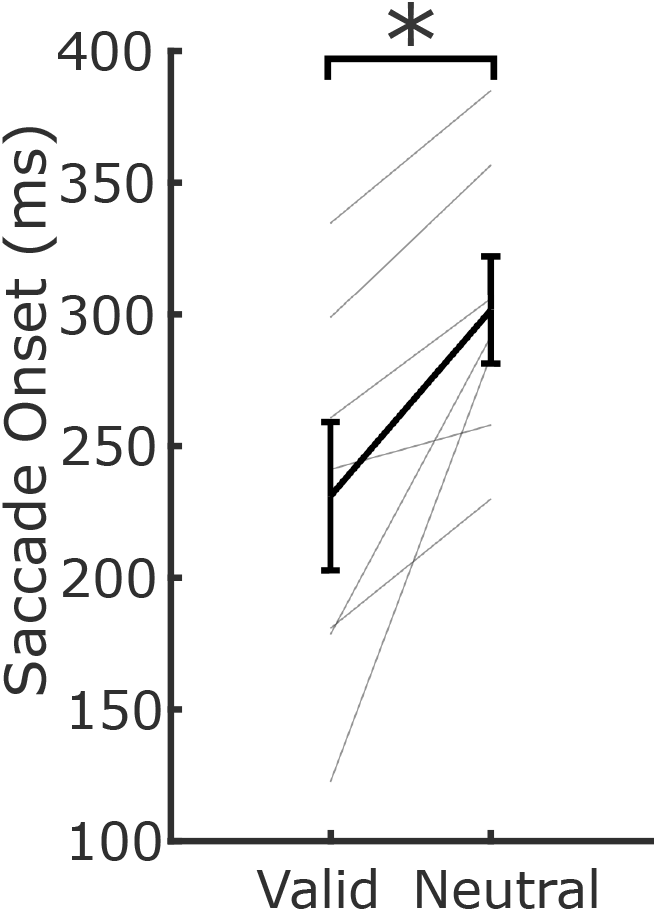
Average saccades onset relative to the response cue onset in valid and neutral conditions (valid vs. neutral: p < 0.01). Only saccades that occurred after fixation period were included in the analysis. The trials presented here were not included in main analyses examining the effects of attention. Lines represent individual observers. Error bars mark group SEM.

**Figure 3—figure supplement 1.**
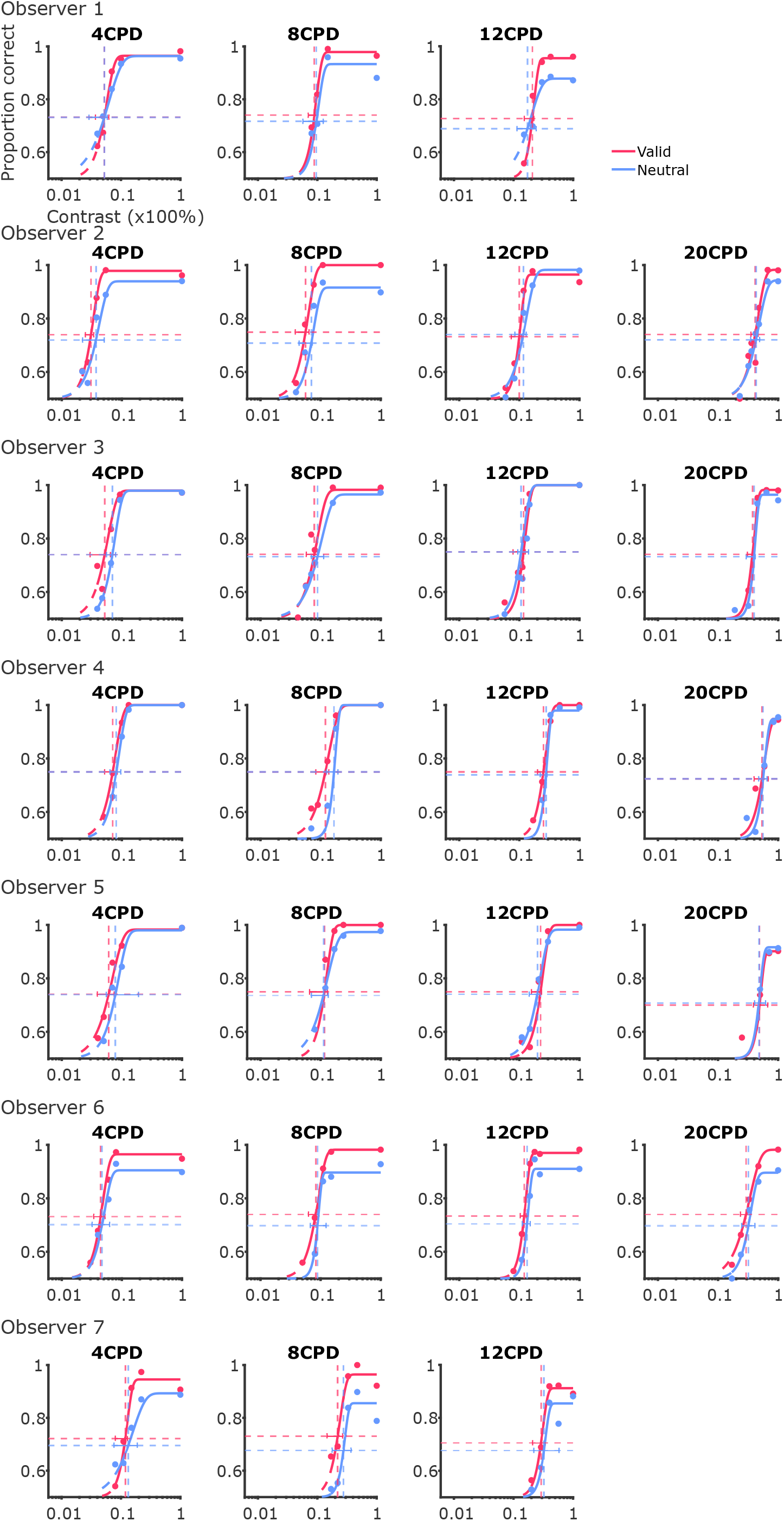
Psychometric Weibull function fits for all observers and conditions.

**Figure 3—figure supplement 2.**
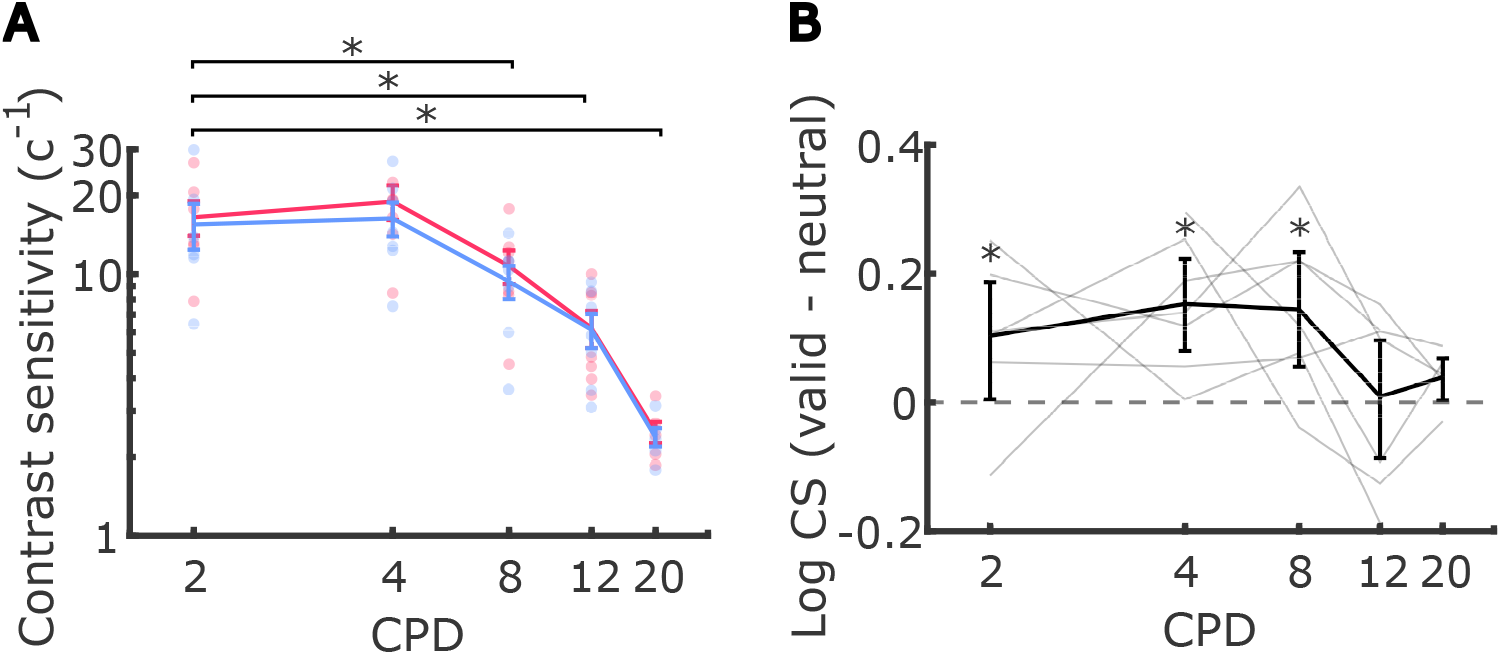
(**A**) Average contrast sensitivity, calculated as the inverse of the contrast threshold, across spatial frequencies, including 2 CPD in valid and neutral conditions. Each dot represents an individual observer. Error bars denote the standard error of the mean (SEM). Asterisks mark a significant difference in contrast sensitivity between 2 CPD and other spatial frequencies. (**B**) Average difference in log-scaled contrast sensitivity between valid and neutral conditions across different spatial frequencies. Each line corresponds to the log-scaled contrast sensitivities from each observer. Error bars represent the bootstrapped 95% confidence intervals. Asterisks mark post-hoc pairwise comparison results between valid and neutral conditions within each spatial frequency (*p* = 0.04 for valid vs. neutral at 2 CPD; *p*s > 0.17 for mean gain at 2 CPD vs. other SFs).

**Figure 3—figure supplement 3.**
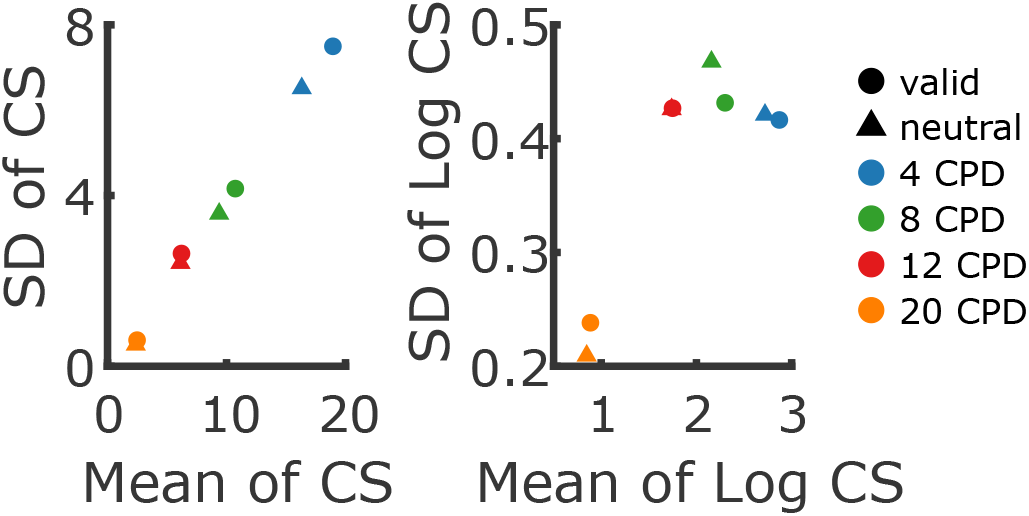
Relation between mean and variability of contrast sensitivity (CS), depending on whether CS is log-transformed (right panel) or not (left panel). Without a log-transform, the mean and standard deviation (SD) of CS are almost perfectly correlated, constituting a strong violation of the homoskedasticity assumption of linear models. For the present data, log-transforming CS *mostly* removes this correlation (except for the 20 CPD condition).

**Figure 3—figure supplement 4.**
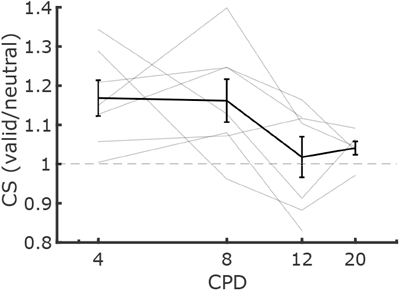
Average attentional benefit calculated as the ratio in contrast sensitivity between valid and neutral conditions.

**Figure 4—figure supplement 1.**
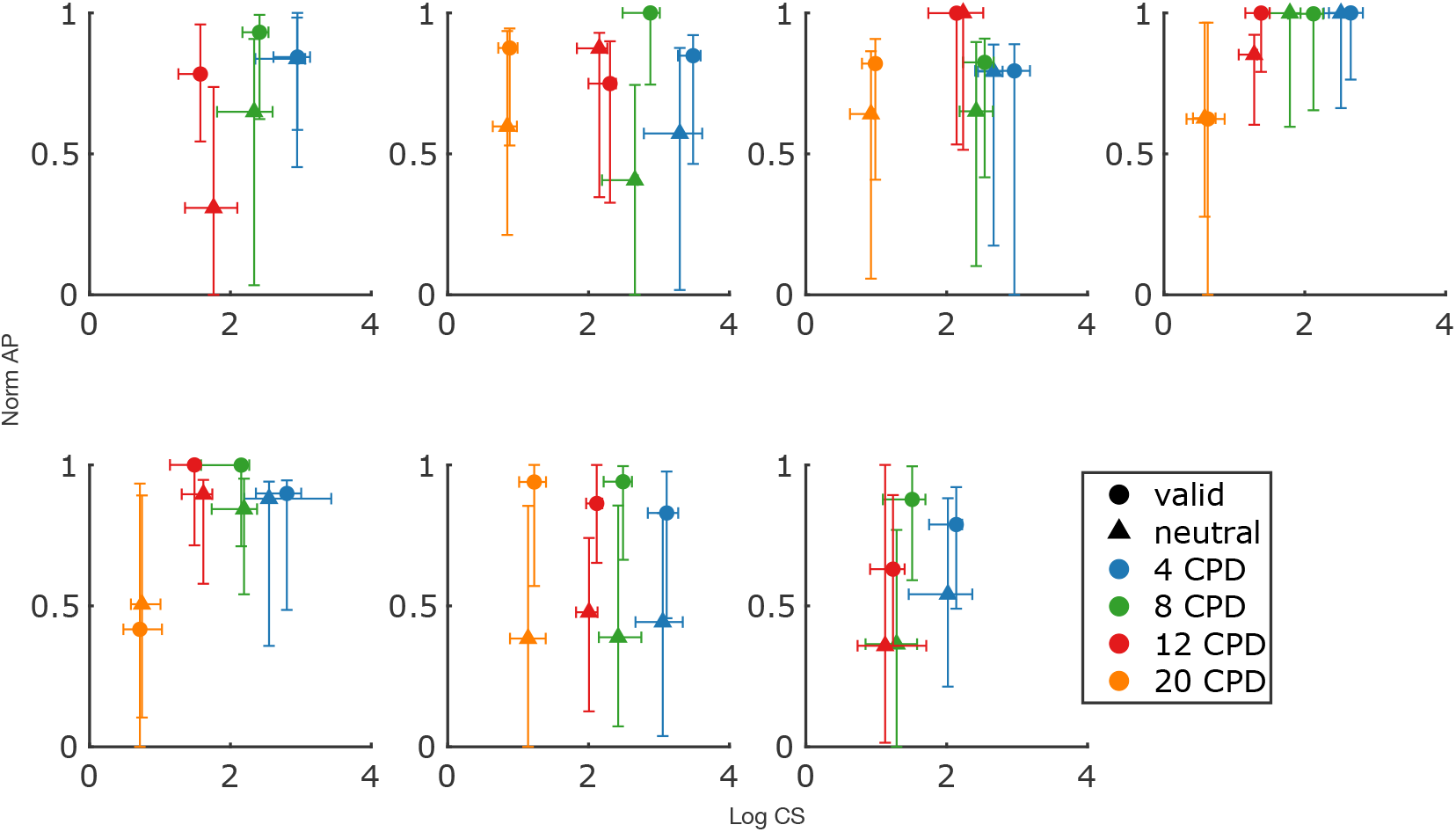
Maximum a posteriori (MAP) estimates (points) and 95% confidence intervals for contrast sensitivity (CS) and asymptotic performance (AP) for each experimental condition across seven observers. Both CS and AP were transformed in the same way as used in our mixed-effects analyses.

## Footnotes

1 Post-hoc pairwise comparisons revealed several significant differences between spatial frequencies (post-hoc pairwise comparisons of the estimated marginal means, *p*s ≤ 0.0001 for 4 vs. 8 CPD, 4 vs. 12 CPD, 4 vs. 20 CPD, 8 vs. 20 CPD, 12 vs. 20 CPD).

2 Post-hoc pairwise comparisons revealed significant differences in asymptotic performance only between 8 and 20 CPD (*p* = 0.015) as well as 12 and 20 CPD (*p* = 0.048).

3 Post-hoc pairwise comparisons between valid and neutral conditions within each spatial frequency revealed that attention significantly increased asymptotic performance compared to neutral condition at 8 and 12 CPD (mean gain_8 CPD_ = 0.05 ±0.04SD, *p* < 0.0001 and mean gain_12 CPD_ = 0.03±0.03SD, *p* = 0.0108) but not at 4 and 20 CPD (mean gain_4 CPD_ = 0.02±0.03SD, *p* = 0.0819 and Δmean gain_20 CPD_ = 0.03±0.04SD, *p* = 0.0501; see ***Appendix 2—Table 2*** for all pairwise comparisons.) When comparing the attentional benefit on asymptotic performance across pairs of spatial frequencies, we found a significant difference (Δmean gain) only between 4 and 8 CPD (*p* = 0.0240; see ***Appendix 2—Table 3*** for all pairwise comparisons).

